# All-optical analysis of electrical coupling in muscle ensembles reveals contributions of individual innexins to cell synchronization and locomotion

**DOI:** 10.64898/2026.03.03.709215

**Authors:** Nora Elvers, Amelie Bergs, Christin Bessel, Jana Liewald, Alexander Gottschalk

## Abstract

Gap junctions (GJs), formed by connexins in vertebrates and innexins in invertebrates, enable direct ion flow between adjacent cells. GJ coupling is typically analyzed by electrophysiological methods, which, however, are invasive and cannot be conducted in intact animals. Here, we used all-optical methods to non-invasively investigate electrical coupling in body wall muscle (BWM) cells of intact *Caenorhabditis elegans*. We analyzed the roles of specific innexins (*unc-9, inx-11, inx-16*) in BWMs, using the genetically encoded fluorescent voltage indicator QuasAr2 to assess spontaneous muscle activity. *unc-9* mutants showed strongly reduced coupling and asynchronous activity, and this was accompanied by severe locomotion defects. Overexpression of the murine connexin Cx36 increased synchronization but disrupted locomotion. *inx-16* mutants exhibited increased excitability of individual muscle cells, likely due to reduced leak currents, while *inx-11* mutants displayed only moderate defects. Patch-clamp recordings confirmed altered action potentials and increased input resistance in *inx-16* mutants. Additionally, we established a cell-specific optogenetic voltage clamp (cOVC) method, directly revealing changes in junctional conductance in live, non-dissected animals. In sum, our findings demonstrate that a balanced level of GJ coupling is essential for coordinated muscle activity and proper motor behavior.

**Significance statement:** Coordination of cells in muscular organs requires electrical coupling *via* intercellular channels termed gap junctions (GJs). How this coupling contributes to coordinated activity like locomotion in an intact animal is difficult to assess, since commonly used electrophysiological methods require dissection, and are incompatible with behavior. Here, we developed all-optical electrophysiology methods to study GJ coupling in live *C. elegans*. Different innexin mutants, as well as overexpression of mammalian connexin Cx36, induced different locomotion defects and, compared to wild type, were accompanied by more or less electrical coupling and synchronicity of individual muscle cells. Loss of some innexins caused the ‘insulated’ muscle cells to be more excitable. Thus, GJ coupling is required for fine-tuning locomotion, and can be studied by voltage imaging.

## Introduction

Gap junctions (GJs) are intercellular channels that provide direct cell-to-cell connections, allowing the exchange of ions, small molecules and metabolites between two adjacent cells. In addition to their important function in the nervous system, where GJs form electrical synapses to enable direct electrical signaling between neurons, GJs are also crucial for the correct propagation of action potentials across the heart and coordinated contraction of intestinal smooth muscles, as well as many other processes (1, 2). GJs are formed from (two) hemichannels, connexons in vertebrates and innexons in invertebrates. Each hemichannel consists of six subunits in vertebrates, the connexins, and six to eight innexins in invertebrates (2–6). Gap junctions can assemble into homo- and heterotypic GJs, whereby the two hemichannels are identical or differ from each other. In addition, each hemichannel can be composed of only one type of connexin/innexin or of multiple different subunits, resulting in homo-or heteromeric GJs, respectively (7). There are 21 different connexin genes in the human genome, while the genome of the nematode *Caenorhabditis elegans* (*C. elegans*) encodes for 25 innexins (8–10). Multiple innexins or connexins are co-expressed in the same cell types, enabling the formation of a large number of different GJs (11–14). Different subunit compositions were shown to affect GJ permeability and gating and thus the intercellular communication they mediate (15, 16). Yet, investigation of GJ composition is challenging, because it is difficult to prove that the different connexins or innexins assemble into the same channel.

Commonly used methods to investigate GJ coupling, like tracer diffusion between cells, fluorescence after photobleaching (FRAP) and local activation of molecular fluorescent probes (LAMP) lack cell-type specificity (17–21). Patch-clamp electrophysiology is highly sensitive and allows the direct measurement of junctional conductance (17, 22, 23). However, electrophysiology is invasive and demanding, especially when used on multiple cells, and in live animals. There are hybrid approaches, where electrophysiology is combined with a voltage-sensitive fluorophore, monitoring network connectivity (24). By combining electrophysiology with dye transfer imaging, heterotypic GJs consisting of mammalian connexins 43 and 45 were investigated with regards to how dye is transferred in response to differences in transjunctional voltage (25). Yet, to study GJ-dependent coupling in a more physiological manner, methods are required that are less invasive, cell-type specific and have a high spatiotemporal resolution. An all-optical approach to measure GJ coupling is PARIS (pairing actuators and receivers to optically isolate GJs), which uses Arch-T, a light-sensitive proton pump, to create an electrochemical gradient, as well as pHluorin, a pH-sensitive variant of green fluorescent protein in the adjacent cell, to detect the movement of protons across GJs into the actuator cell (26). However, proton transfer is rather artificial, thus it would be beneficial to use imaging to analyze native ion fluxes in live animals.

Optical readout of changes in membrane potential can be achieved by using genetically encoded voltage indicators (GEVIs). Rhodopsin-based GEVIs show a voltage-dependence of the fluorescence of their retinal chromophore, enabling the detection of electrical activity at millisecond timescales (27–34). Since rhodopsin-based GEVIs emit near-infrared light, they can be combined with optogenetic actuators (28, 35–37). The recently established optogenetic voltage clamp (OVC) allows the clamping of cells to a specific holding potential by using the GEVI QuasAr2 for voltage read-out, and the actuator protein BiPOLES, consisting of the depolarizer Chrimson and the hyperpolarizer *Gt*ACR2 (38). This system enables the clamping of voltage in muscular organs and neurons in *C. elegans*, and could detect altered cell physiology in mutants.

The 95 spindle-shaped body wall muscle (BWM) cells of *C. elegans* are organized into four quadrants that run along the length of the body and allow the animals to move in a coordinated sinusoidal movement. *Via* GJs in muscle arms, BWM cells are connected to their nearest neighbors on the dorsal or ventral side, allowing inter-quadrant coupling. Moreover, within a quadrant, muscle cells are connected to their immediate neighbors via GJ plaques along the connecting cell-cell border, i.e. where their plasma membranes are in direct contact (23, 39–42). The muscle cells receive input from cholinergic and GABAergic motor neurons, leading to coordinated contraction and relaxation of the cells, thus mediating the propagation of the undulatory wave (43). Up to now, the function of GJ-mediated coupling between the muscle cells is unclear. In the absence of neurotransmission, muscle cells are still able to generate action potentials (APs) and Ca^2+^ transients, indicating intrinsic muscle activity (44). Inhibition of acetylcholine and GABA transmission led to decreased AP frequencies and Ca^2+^ transients, however, APs and Ca^2+^ transients remained synchronous, indicating that GJs might facilitate synchronization of muscle cells (42). Using electrophysiological recording, six innexins were shown to be involved in the electrical coupling between muscle cells. Based on these experiments, Liu et al. proposed that two populations of heterotypic GJs are involved, one consisting of UNC-9 and INX-18, and the other of INX-1, INX-10, INX-11 and INX-16 (12).

Here we show that GJ-mediated electrical coupling can be analyzed using genetically encoded voltage indicators. We use the GEVI QuasAr2 (28) to investigate the GJ-dependent coupling of BWMs. Effects on the correlation of intrinsic BWM activity were observed for *unc-9* mutants, confirming a major role of UNC-9 in electrical signaling in BWMs. Mutations of INX-11 and INX-16 had only minor effects on locomotion and the coordination of BWMs. However, in *inx-16* mutants, we could show that single muscle cells were more isolated from each other, meaning less currents leak to neighboring cells, leading to increased excitability of individual cells. Furthermore, we expanded the previously established optogenetic voltage clamp (38) to enable optical clamping of one individual cell and to read out the GJ-mediated voltage changes in the neighboring cells. Using this approach, we were able to show an overall lower electrical conductivity in *unc-9* mutants and a higher excitability of individual cells in *inx-11* mutants. Furthermore, we show that overexpression of a heterologous GJ, consisting of the murine connexin Cx36, evokes locomotion defects due to an overall higher muscle cell correlation, emphasizing that the right balance of GJs expression is required for the BWM network to function properly.

## Results

### Loss of innexins as well as increased GJ coupling lead to different locomotion phenotypes

In order to study GJ-mediated electrical coupling, most often patch-clamp electrophysiology is used, as it enables the direct measurement of junctional conductance. However, it is technically demanding and requires dissection of the animals, and thus cannot be used in live, behaving animals, complicating the interpretation of the results (17, 22, 23). In contrast, genetically encoded voltage indicators (GEVIs) report on millisecond dynamics of electrical signaling and can be applied to study voltage dynamics of neurons or muscles in a spatiotemporal manner in live animals, without the need for dissection (27, 28, 36, 45). Mutants of the innexin-encoding genes *unc-9, inx-16* and *inx-11* showed the strongest effects in junctional conductance compared to WT animals (12). Based on these previous findings, we chose these three innexins to establish the application of voltage imaging techniques for the analysis of cellular coupling in muscular ensembles.

We first assessed locomotion behavior, since reduced GJ coupling was shown to affect muscle action potential synchrony and locomotion (23, 42). While wild type (WT) animals showed 116.5±1.6 µm/s average crawling speed in the absence of food (Fig. 1A-B), *unc-9* mutants were almost immotile (20.9±0.8 µm/s), which is in accordance with previous studies (23).

**Figure 1:**
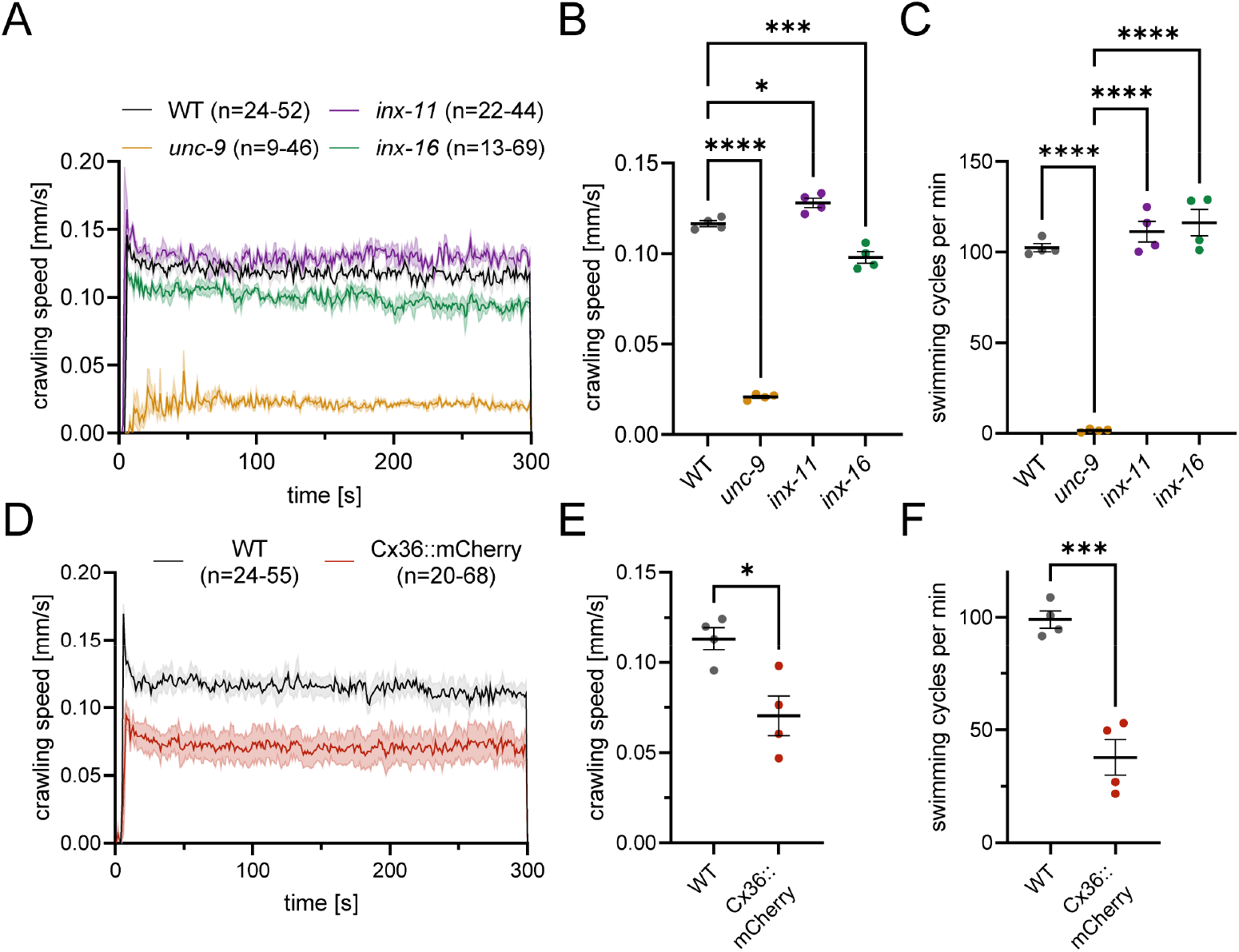
Loss of innexins or increased GJ levels lead to opposing effects on locomotion. (A) Basal crawling speed (±SEM) of *unc-9(e101), inx-11(ok2783)* and *inx-16(tm1589)* mutants compared to WT animals. Number of animals per experiment n, across N=4 independent experiments. (B) Comparison of mean crawling speed of the indicated strains. Each dot represents the mean crawling speed of each measurement. Mean and SEM. One-way ANOVA with Dunnett test (* p < 0.05, *** p < 0.001, **** p < 0.0001). (C) Mean swimming cycles (±SEM) of WT, *unc-9, inx-11* and *inx-16* mutants. Mean and SEM. Number of animals per experiment n=24-52 (WT), n=34-86 (*unc-9*), n=28-51 (*inx-11*) and n=18-72 (*inx-16*), across N=4 independent experiments. One-way ANOVA with Tukey test (**** p < 0.0001). (D) Basal crawling speed (±SEM) of Cx36::mCherry overexpressing animals compared to WT. Number of animals per experiment n, across N=4 independent experiments. (E) Comparison of mean crawling speed of the indicated strains. Each dot represents the mean crawling speed of each measurement. Mean and SEM. Unpaired t test (* p < 0.05). (F) Mean swimming cycles (±SEM) of WT and Cx36::mCherry overexpressing animals. Number of animals per experiment n=32-68 (WT) and n=26-74 (Cx36), across N=4 independent experiments. Unpaired t-test (*** p < 0.001).

Mutants lacking the innexin INX-16 were significantly slower than WT (98±3 mm/s) and *inx-11* mutants were slightly but significantly faster (128.0±3.0 mm/s). We analyzed a second locomotion gait, i.e. swimming, which showed strong defects in *unc-9* mutants, while *inx-11* and *inx-16* mutants were unaffected (Fig. 1C), indicating that different innexins influence aspects of locomotion differently, and further, that crawling and swimming behaviors are differently affected by loss of individual GJ subunits.

We also assessed a gain-of-function situation. To this end, we overexpressed a heterologous GJ, the murine connexin Cx36, in BWMs. Cx36 forms homotypic/homomeric GJs and was previously shown to be functional in *C. elegans*, presumably forming homotypic GJs with other Cx36 molecules, but is not expected to form GJs with innexins endogenous to *C. elegans* (46, 47). Cx36::mCherry, expressed from the *myo-3* promotor, showed punctate expression along the plasma membrane of BWMs (Supp. Fig. 1A). Overexpression of Cx36::mCherry resulted in overall lower crawling and swimming speed compared to WT animals (Fig. 1D-F), indicating that overexpression was negatively affecting muscular coupling, or, alternatively, was toxic to muscle cells. However, the latter appears unlikely, given the normal shape of BWMs (Supp. Fig. 1A).

### Voltage imaging reveals asynchronous muscle activity in *unc-9* mutants, while overexpression of Cx36 increased coordination

Next, we analyzed how these innexin mutants or overexpression animals, i.e. reduction / loss-of-function, as well as gain-of-function affected electrical coupling. To this end, we used the GEVI QuasAr2 (27, 28, 38). Expression of QuasAr2 in BWMs allowed us to measure voltage-dependent fluorescence fluctuations, which are likely evoked by action potentials (APs), or bursts of APs (44). We assessed ensembles of three neighboring muscle cells in the midbody region, adjacent to the vulva, where the longitudinal overlap of cells 1, 2 and 1, 3 was large, while cells 2, 3 had less direct overlap (Fig. 2A; Supp. Movie 1). Analyzing fluorescence fluctuations in WT animals (Fig. 2B) showed highly similar activity in all three cells. In line with previously reported calcium imaging data (42), *unc-9* mutants exhibited asynchronous muscle activity during voltage imaging, as well as an overall reduced correlation (measured over the entire recording period) in comparison to WT (Fig. 2C, E; Supp. Fig. 1B). The characteristics of the peaks, like amplitude, area under the curve and full width at half maximum (FWHM) did not differ from WT (Fig. 2F-H). In contrast, Cx36 overexpressing muscles acted much more correlated than the WT (Fig. 2B, D, I; all three WT controls showed similar overall correlation of muscle AP activity; Supp. Fig. 1C). Single AP-evoked fluorescence peaks did not differ from WT, though the peaks were slightly, but significantly narrower (Fig. 2J-L), possibly reflecting more precise and acute signal propagation between muscle cells. This may preclude the normal propagation of the undulatory wave of muscle contractions associated with *C. elegans* locomotion. Thus, the locomotion phenotype of Cx36 overexpressing animals is likely due to excessive BWM coupling.

**Figure 2:**
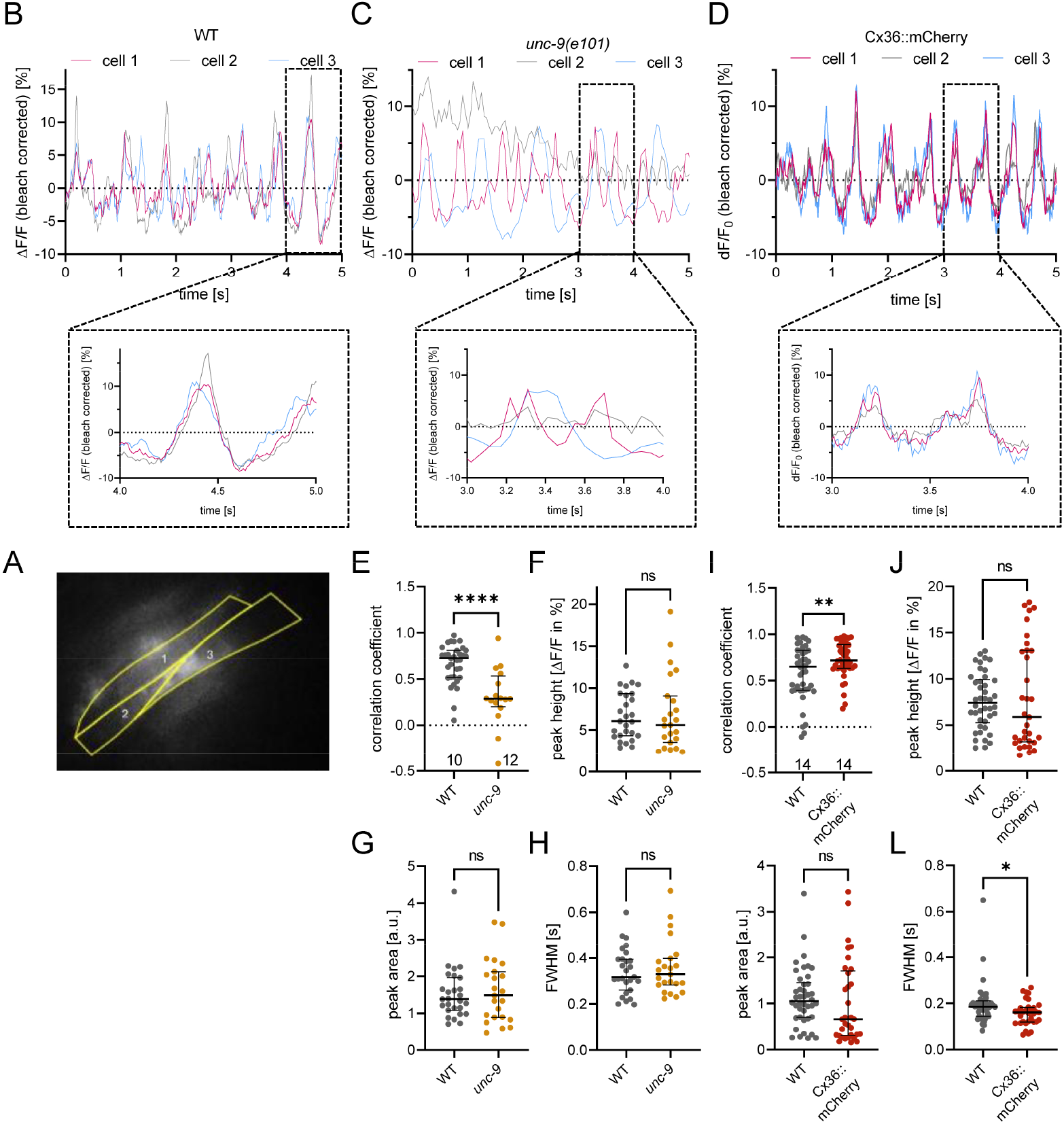
Spontaneous muscle cell activity is visualized by QuasAr2 imaging, revealing differences in *unc-9* mutants and Cx36::mCherry overexpressing muscles. (A) Fluorescence of QuasAr2 expressed in body wall muscles of *C. elegans*. Measured were ensembles consisting of three muscle cells, yellow outline shows the analyzed ROIs. Numbering of cells is indicated. (B) Representative ΔF/F fluorescence time traces of spontaneous muscle cell activity of WT animals, *unc-9* mutants (C) and Cx36::mCherry overexpressing animals (D). Correlation of muscular activity WT, *unc-9* mutants (E) and Cx36::mCherry overexpressing animals (I) was calculated using Pearson correlation. Each dot represents one calculated correlation coefficient. Median with interquartile range. Number of animals n=10 (WT, *unc-9*); n=14 (WT, Cx36). Unpaired t test (**** p < 0.0001), Welch’s t test in I. Single peaks of voltage fluorescence traces were analyzed for amplitude (F, J), peak area (G, K) and full width at half maximum (FWHM) (H, L). Each dot represents the mean result of all peaks of one fluorescence time trace of one animal. Median with interquartile range. Number of animals is the same as in E and I, respectively. Mann-Whitney test in F, K, L; unpaired t test in G, H, J (ns p>0.05).

We analyzed the electrical correlation of individual muscle cells in more detail, to see if coordination was affected by the length of cell contacts in the three pairs of cells (1-2, 1-3, 2-3; Fig. 1A). For WT, we found that the cell pair 2-3, with the smaller overlap, showed less correlation (Supp. Fig. 1D). For Cx36 expressing animals, we speculated that the presence of the heterologous connexin might abolish these differences, because of its abundance and putatively random localization. Indeed in Cx36 animals, all cell pairs were equally correlated, and cell pair 2-3 showed significantly better correlation than in WT animals (Supp. Fig. 1D). For other strains / mutants, this did not show any obvious differences, so these data were pooled. Cross correlation analysis of single voltage fluorescence peaks (± 0.25 s) from the three analyzed neighboring muscle cells were performed using a Matlab script. Recordings from *unc-9* mutants, compared to WT, showed significantly decreased maximal correlation coefficients among muscles (Fig. 3A, B; see Supp. Fig. 2A for mean fluorescence traces). No difference was observed between individual muscle pairs in WT. This was also the case for cross correlation of fluorescence traces during longer time windows, centered on a central peak, ± 0.65 s, and sometimes containing additional peaks (Supp. Fig. 2B-E), and confirmed that *unc-9* mutants suffer from uncoordinated activity in adjacent muscles. The distribution of the lag times at maximal correlation coefficient was broader in *unc-9* mutants, however the mean lag times were not significantly different from WT (Fig. 3C), again suggesting that APs occurred in a highly unsynchronized manner in neighboring muscle cells of *unc-9* mutants, likely due to reduced electrical coupling. Normalization of the cross correlation of extracted peaks illustrates the asynchronous muscle activity in *unc-9* mutants (Fig. 3D). Fitting of the decay was only possible for the cross correlation of cell pair 1-2 yielding a τoff of 0.098 s, which, compared to WT (τ_off_=0.21±0.04 s), exhibited a faster decay and reduced coupling over time. For Cx36 overexpressing muscles, cross correlation analysis revealed a higher maximal correlation coefficient, without affecting lag time (Fig. 3H-J, Supp. Fig. 2F), further demonstrating increased coordination of adjacent muscles. Tau_off_ of the normalized cross correlation of adjacent Cx36 muscle cells was not significantly different from WT (Fig. 3K, L). Cross correlation of a longer time window revealed a higher maximal correlation coefficient compared to WT, indicating again more precisely coordinated activity of the three muscle cells (Supp. Fig. 2G-J). These effects were very similar for individual muscle pairs.

**Figure 3:**
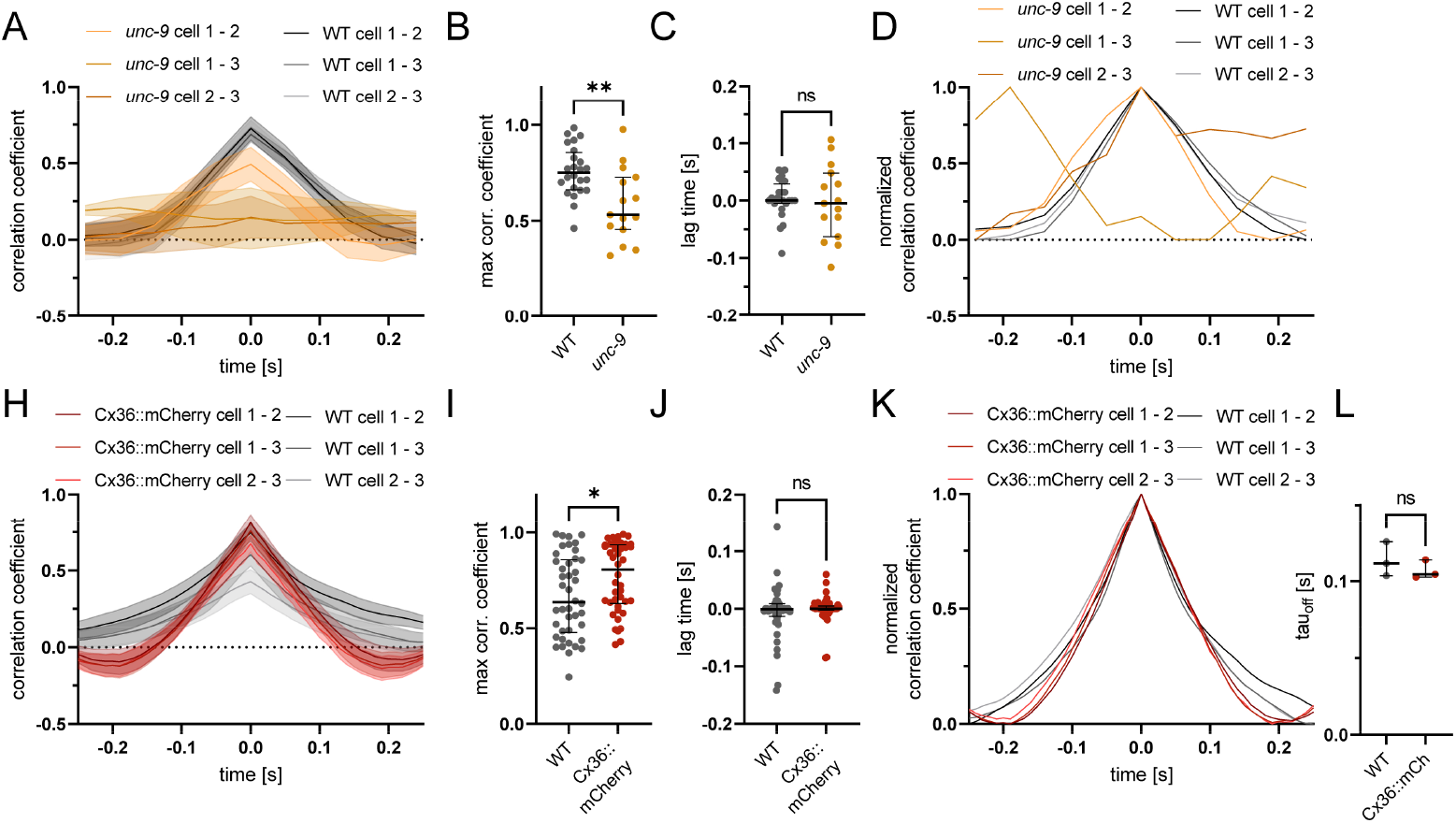
Cross correlation analysis of single voltage fluorescence peaks reveals changes in maximum correlation. (A) Cross correlation of single peaks (± 0.25 s from peak) of pairs of the three-cell ensemble of *unc-9* mutants and WT. Mean and SEM. Number of animals n=9-10 (WT), n=3-9 (*unc-9*). Maximum correlation coefficient (B) and lag time at maximum correlation (C) of the cross correlations of ten peaks from each animal. Each dot represents the mean of the first 10 cross correlations of one pair of muscle cells. Median with interquartile range. Unpaired t test (** p<0.01) in B, Welch’s t test (ns p<0.05) in C. (D) Min-max normalization of the cross correlation of single peaks in A. (H) Cross-correlation analysis of single peaks (±0.25 from peaks) of pairs of the three-cell ensemble of Cx36::mCherry overexpressing animals. Number of animals n=14 (WT, Cx36). (I) Maximum correlation coefficient and lag time at maximum correlation (J) of the cross correlation of the ten peaks from each animal. Each dot represents the mean of the first 10 cross correlations of one pair of muscle cells. Median with interquartile range. Mann-Whitney test (ns p>0.05, * p<0.05). (K) Min-max normalization of the mean cross correlations shown in H. (L) Calculated tau_off_ value of the mean cross correlation of a pair of muscle cells using a one-phase exponential decay fit. Median with interquartile range. Unpaired t test (ns p>0.05).

### Voltage imaging of spontaneous muscle activity indicates higher excitability in *inx-16* mutants

To show how effective voltage imaging is for investigating presumably small alterations in electrical signaling, we analyzed *inx-11* and *inx-16* mutants that had shown minor alterations in locomotion behavior, compared to wild type (see Fig. 1). Spontaneous BWM activity resembled WT recordings (Fig. 4A-C), showing in some cases that the voltage fluorescence peaks actually consist of bursts of several action potentials (APs). These occurred roughly at a frequency of 3 Hz and contained several (3–5) discernible APs. The temporal resolution of voltage imaging is determined by the frame rate of the camera and the achievable photon count for very brief exposure times (38). When we compared the coordination of activity in the three cells, this showed no difference between WT, *inx-11*, and *inx-16* mutants in overall correlation (Fig. 4D). Also, cross correlation analysis of single voltage peaks of *inx-11* mutants revealed no differences compared to WT animals (Fig. 5A, C-E, G; Supp. Fig. 2K, M, O, Q, R). However, *inx-16* mutants showed slightly increased peak amplitudes and significantly larger peak area and FWHM compared to WT (Fig. 4E-G), indicating longer AP (bursts). This could suggest that single muscle cells are more isolated in the absence of INX-16(-containing GJs), since less current may be leaking into the neighboring cell. This is expected to increase the excitability of the individual BWM cell and voltage rises in *inx-16* mutants. Since *inx-16* mutants showed slightly impaired crawling, this suggests that this increased isolation leads to reduced smoothness in locomotion wave propagation. Cross correlation analysis of single voltage peaks / bursts of APs (± 0.25 s from peak), between mutual pairs of individual muscle cells of the respective three-cell ensemble, showed a significantly increased maximal correlation coefficient of *inx-16* mutants (Fig. 5B-C, Supp. Fig. 2L), showing increased synchronicity of neighboring muscle cells, while the lag time at maximal correlation was unaffected (Fig. 5D). The cross correlation tau_off_ value was significantly longer in *inx-16* mutants than in WT or *inx-11* mutants (Fig. 5E-G), due to overall broader AP bursts. We also looked at cross correlations of longer time periods (± 0.65 s from peak), where no differences in correlation coefficients or lag times were detected (Supp. Fig. 2N, P, Q, R). Together, these findings demonstrate that specific innexin subunits affect the electrical coupling of muscle cells through GJs with respect to ensemble activity but also AP (burst) characteristics. Using voltage imaging, changes in GJ mediated muscle coupling can be analyzed in a mostly physiological manner, by assessing intrinsic, spontaneous muscle cell activity in intact animals.

**Figure 4:**
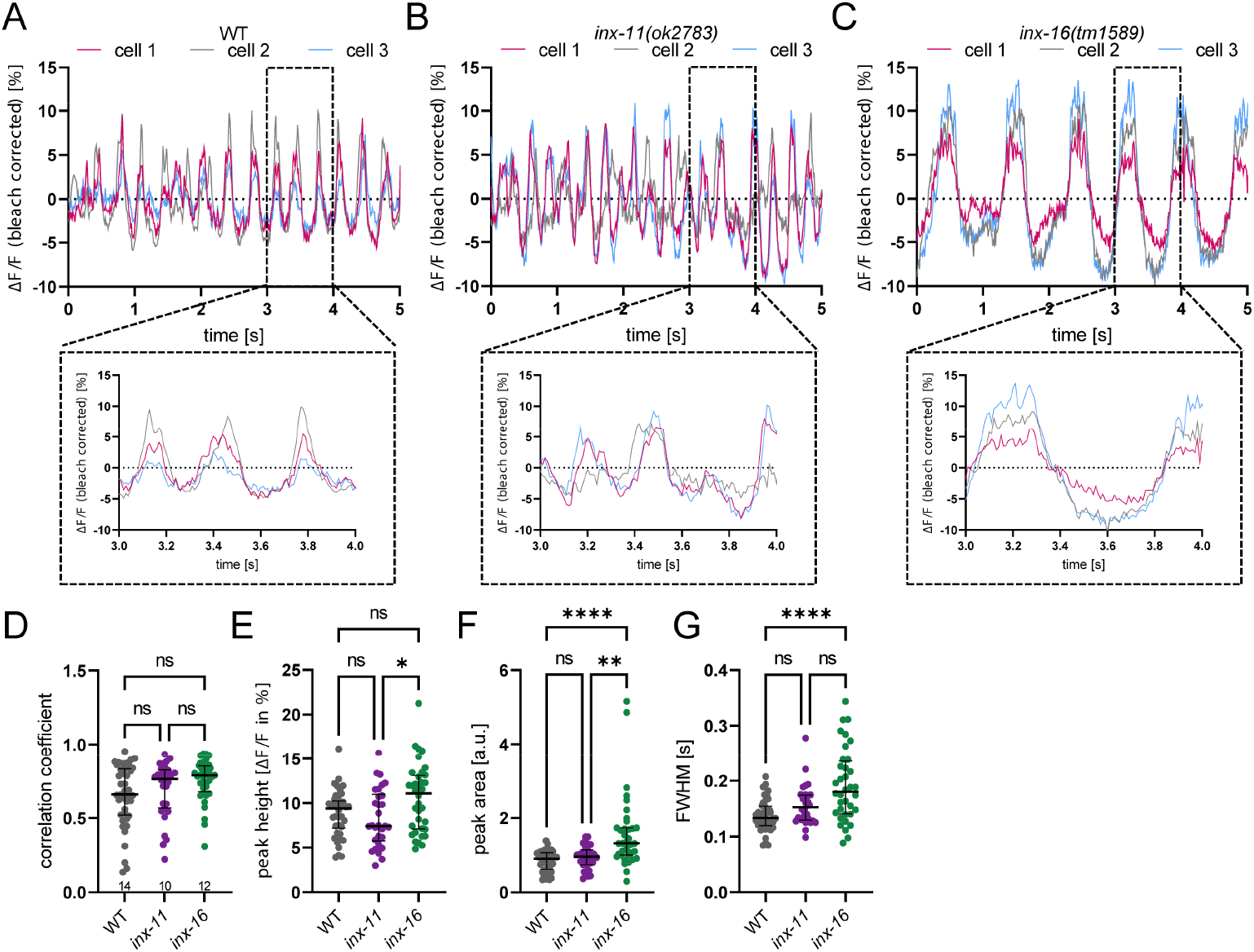
Voltage imaging of spontaneous muscle activity reveals changes in peak characteristics in *inx-16* mutants. (A) Representative traces of spontaneous muscle cell activity of one WT animal. Representative ΔF/F fluorescence time traces of *inx-11* mutants (B) and *inx-16* mutants (C). (D) Correlation of muscular activity was calculated using Pearson correlation. Each dot represents one calculated correlation coefficient. Median with interquartile range. Number of animals n=14 (WT), 10 (*inx-11*), 12 (*inx-16*). Kruskal-Wallis with Dunn’s test (ns p>0.05). Single peaks of voltage fluorescence traces were analyzed for amplitude (E), peak area (F) and FWHM (G). Each dot represents the mean result of all peaks of one fluorescence time trace of one animal. Median with interquartile range. Number of animals as in D. One-way ANOVA with Tukey test in E, Kruskal-Wallis with Dunn’s test in F, G (ns p>0.05, * p<0.05, ** p<0.01, **** p<0.0001).

**Figure 5:**
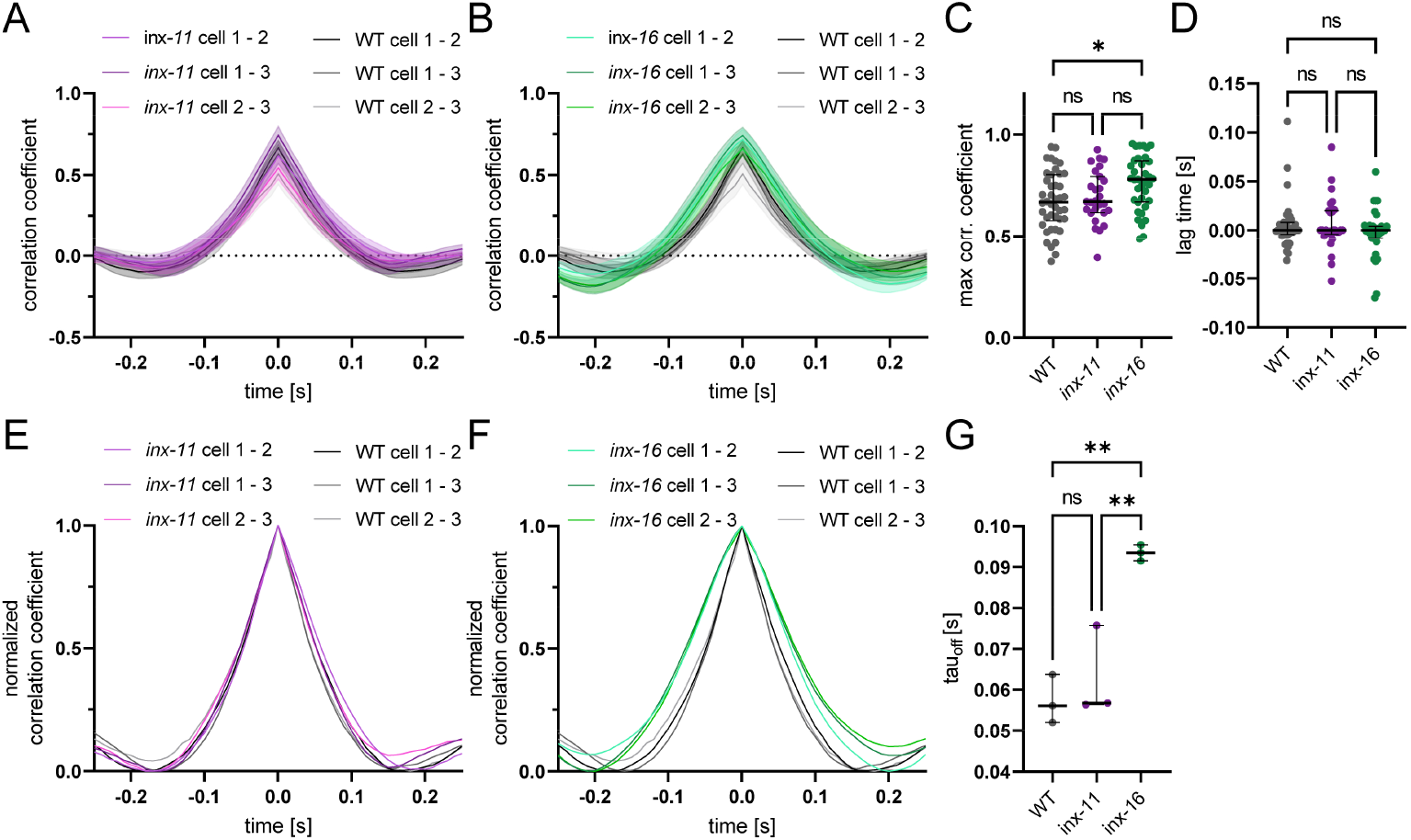
Cross correlation analysis of single peaks reveals higher maximum correlation of *inx-16* mutants. Cross correlation of single peaks (± 0.25 s from peak) of pairs of the three-cell ensemble of *inx-11* (A) and *inx-16* mutants (B). Mean and SEM. Number of animals n=14 (WT), 9 (*inx-11*), 12 (*inx-16*). Maximum correlation coefficient (C) and lag time at maximum correlation (D) of the cross correlations of ten peaks from each animals. Each dot represents the mean of the first 10 cross correlations of one pair of muscle cells. Median with interquartile range. One-way ANOVA with Tukey test (ns p>0.05, * p<0.05) in C, Kruskal-Wallis with Dunn’s test (ns p<0.05) in D. (E) Min-max normalization of the cross correlation of single peaks in A for *inx-11*, and in B for *inx-16* mutants. (G) Calculated tau_off_ value of the mean cross correlation of a pair of muscle cells using a one-phase exponential decay fit. Median with interquartile range. One-way ANOVA with Tukey test (ns p>0.05, ** p<0.01).

### Patch clamp recordings indicate increased excitability of *inx-16* mutants, in line with GEVI measurements

Our findings in voltage imaging of adjacent muscles indicated that mutations of innexins, as well as heterologous (and possibly, ectopic) GJ expression, can affect the electrical coupling in opposing ways, leading to more or less coupling, and affect AP burst characteristics (*inx-16*). To directly assess the electrophysiological characteristics and excitability of muscle cells, we conducted patch clamp recordings of BWM cells in dissected animals. Though dissection of the worm and the use of artificial buffers may alter the native spontaneous activity of muscles in electrophysiology recordings, general cell physiological parameters can be precisely measured and compared. No significant differences in resting membrane potential were found in innexin mutants compared to WT, though *unc-9* mutants showed a larger mean value (Fig. 6A). Mutants of *inx-16* showed a significant decrease of membrane capacitance compared to WT (Fig. 6B), indicating a smaller cell size. The reduced size was obvious by eye, in line with a previous report stating that *inx-16* mutants are smaller and grow more slowly (48). Cellular input resistance, normalized to the capacitance, was increased in *inx-16* mutants, indicating a higher excitability of the muscle cells (Fig. 6C).

**Figure 6:**
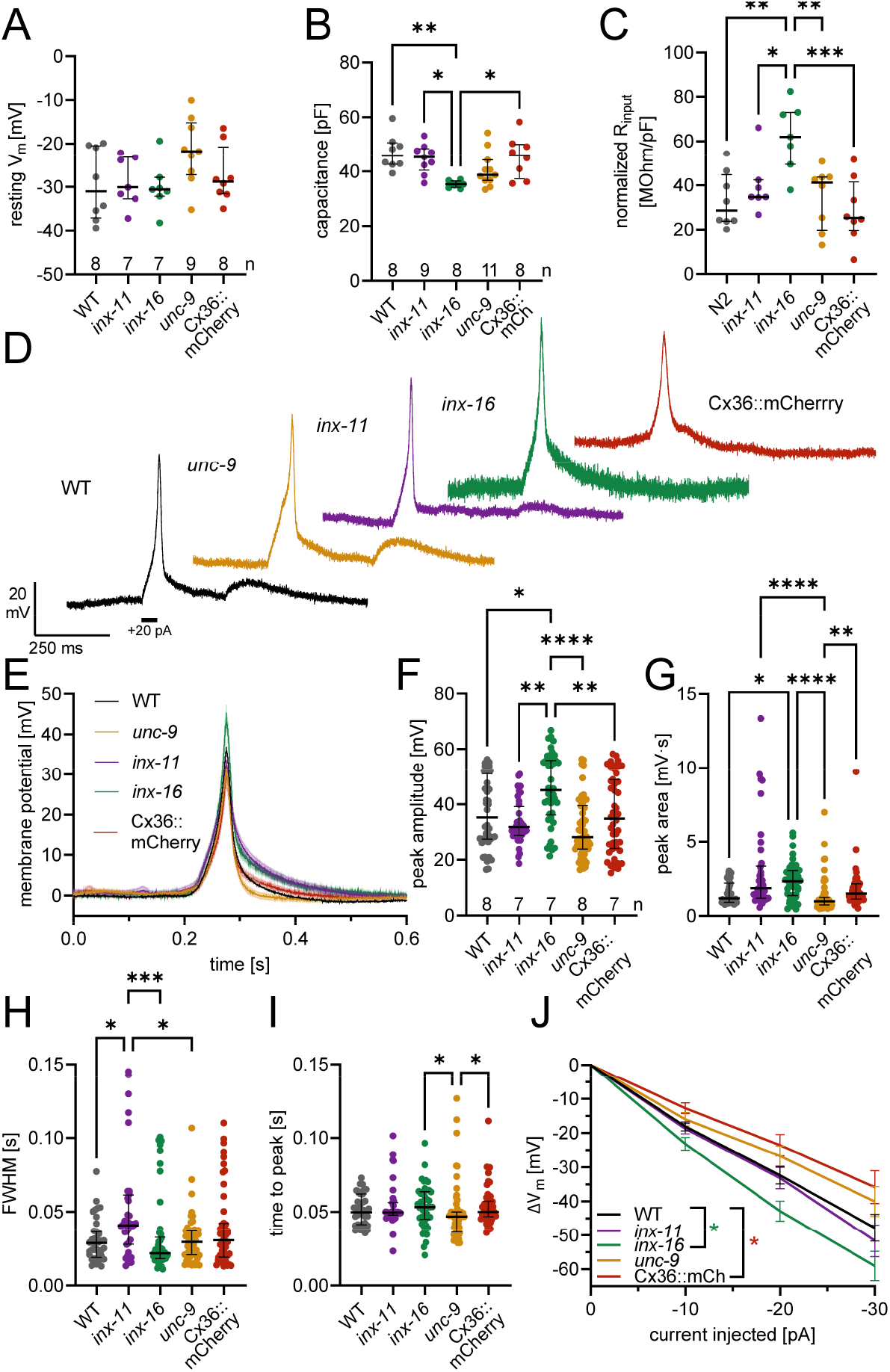
Patch clamp recordings reveal increased excitability of *inx-16* mutant muscles. (A) Resting membrane potential recorded in current clamp of WT, *inx-11, inx-16, unc-9* mutants and Cx36::mCherry overexpressing muscles. Median with interquartile range. Number of animals n=8 (WT), 7 (*inx-11, inx-16*), 9 (*unc-9*), 8 (Cx36). One-way ANOVA with Tukey test. (B) Capacitance measurements in current clamp mode. Median and interquartile range. One-way ANOVA with Tukey test (* p<0.05, ** p<0.01). Number of animals n=8 (WT, *inx-16*, Cx36), 9 (*inx-11*), 11 (*unc-9*). (C) Input resistance was measured by injection of three consecutive −20 pA pulses (1000ms, 1000ms interval). For normalization, the input resistance was divided by the capacitance of the respective cell. Number of animals n=8 (WT, *unc-9*, Cx36), 7 (*inx-11, inx-16*). Median with interquartile range. One-way ANOVA with Tukey test (* p<0.05, ** p<0.01, *** p<0.001). (D) Exemplary traces of induced APs. APs were induced by a 50 ms, +20 pA current pulse. (E) The induced APs were extracted and peak aligned (mean ± SEM). Number of extracted peaks n=40 (WT, *inx-11*, Cx36), 54 (*unc-9*), 46 (*inx-16*). APs were analyzed for amplitude (F), area under the curve (G), FWHM (H) and time to peak (I). Median with interquartile range. Kruskal-Wallis with Dunn’s test (* p<0.05), ** p<0.01, *** p<0.001, **** p<0.0001). (J) Changes of membrane potential in response to current injections of −10, −20 and −30 pA pulses (1000 ms). Number of animals n=8 (WT, *unc-9*, Cx36), 7 (*inx-11, inx-16*). Mean and SEM. Two-way ANOVA with Tukey test (* p<0.05).

Next, to assess excitability more directly, we used brief (50 ms) depolarizing current injections of 20 pA, in order to evoke APs in these cells, and to analyze their shape, duration, and timing (Fig. 6D). Single evoked APs showed canonical shape and duration. We analyzed all induced APs following baseline correction and peak alignment, using an automated Matlab script. The mean APs demonstrated a mostly uniform rising phase for all genotypes tested, however, peak amplitudes and, particularly, duration and voltage change rates during repolarization were slightly different among genotypes (Fig. 6E). While there was only a slightly slower decline for Cx36 overexpressing cells compared to WT (τ_off_=0.053 s (WT), 0.072 s (Cx36)), *inx-16* and *inx-11* mutants showed a slower decline (τ_off_=0.086 s (*inx-11*) and 0.098 s (*inx-16*)), while a faster decline was seen for *unc-9* mutants (τ_off_=0.026 s) (Fig. 6E). Peak amplitude and area were increased for *inx-16* mutants, which is in accordance with the voltage imaging results (Fig. 6F, G), and confirmed that reduced coupling to neighboring cells may increase excitability, as less leak currents leave the individual muscle cell. Why this is not the case for *unc-9* mutants, in which coupling should be reduced even more, is unclear. The discrepancy between the *unc-9* and *inx-16* mutants may further depend on the remaining or reorganized GJs, in the absence of the respective innexin, which may have different functional properties (e.g., conductance, closing kinetics, or rectification). Mutants of *inx-11* showed a significantly increased FWHM compared to WT, in line with broader APs (Fig. 6H). We also analyzed the time to peak from the onset of the current injection, which showed no major differences (Fig. 6I). Last, we compared the change in membrane voltage in response to hyperpolarizing current injections (to prevent AP generation) of increasing amplitude (−10, −20, −30 pA; Fig. 6J). In these experiments, *inx-16* mutants showed significantly larger changes in membrane potential than WT, while Cx36 overexpression led to significantly reduced voltage changes, in line with the idea that more current leaves the patch-clamped Cx36 expressing cell through GJs, and that *inx-16* cells are more insulated.

### Establishment of a cell-specific optogenetic voltage clamp (cOVC)

To directly assess junctional coupling in live, intact animals, we sought to develop an all-optical approach. We previously established the optogenetic voltage clamp (OVC) (38). This method uses optogenetic tools for hyperpolarization and depolarization (BiPOLES, consisting of Chrimson, a Na^+^ channel activated by 590 nm light, and *Gt*ACR2, a Cl^−^ channel activated by 470 nm; (49)), as well as fluorescence based voltage imaging using QuasAr2, to control the membrane potential of excitable cells. To this end, QuasAr2 fluorescence is sampled and this information is used to compute a feedback of wavelength-adapted light to BiPOLES, thus enabling closed-loop control of the membrane potential (38). Now, to investigate the coupling between adjacent cells, we adapted the OVC to allow clamping one muscle cell (i.e., by current pulses injected into this cell), while reading out changes of voltage in the adjacent cell (Fig. 7A; cell-specific OVC - cOVC). Earlier, BiPOLES excitation light was applied to all cells in the field of view. We now exchanged the previously used monochromator with a video projector, allowing to project shapes adjusted to individual muscle cells, in different colors of different intensities (Supp. Fig. 3A). Whereas the original OVC approach relies on wavelength modulation, the present implementation clamps the membrane potential via intensity modulation of two fixed wavelength bands (blue and green). The spectral output of the beamer was narrowed using filters (Supp. Fig. 3B), and the accuracy of illuminated and measured ROIs / pixel size was characterized, as well as the intensities of the different wavelengths (blue: 450-477 nm, green: 552-587 nm) were compared to guarantee robust and ongoing BiPOLES activation (Supp. Fig. 3C-E). A custom-written Beanshell-script was used to define a region of interest (ROI) covering one cell, and to illuminate this cell with the intensity-adapted light, determined by the measured voltage-dependent relative changes in fluorescence of this cell, as compared to the target value. This way, it was possible, to clamp this cell to a desired membrane potential. A second ROI covered the neighboring cell to monitor changes of membrane potential only (Fig. 7B; Supp. Movie 2). To exclude that BiPOLES molecules expressed in cell 2 are activated by scattered light targeted at cell 1, we set one ROI outside the worm, adjacent to a muscle cell whose fluorescence changes were imaged and analyzed. No relative fluorescence changes were observed between the two ROIs (Supp. Fig. 3F), indicating that the observed changes of cell 2 were due to coupling of the neighboring muscle cells and not due to stray light. Control measurements of a strain expressing only QuasAr2 in BWMs showed no effect of the light at wavelengths used for BiPOLES activation on QuasAr2 fluorescence (Supp. Fig. 3G). With this system it was also possible to clamp two cells in two opposing directions (Supp. Fig. 3H), and clamping of single cells could be sustained for dozens of seconds (Supp. Fig. 3I).

**Figure 7:**
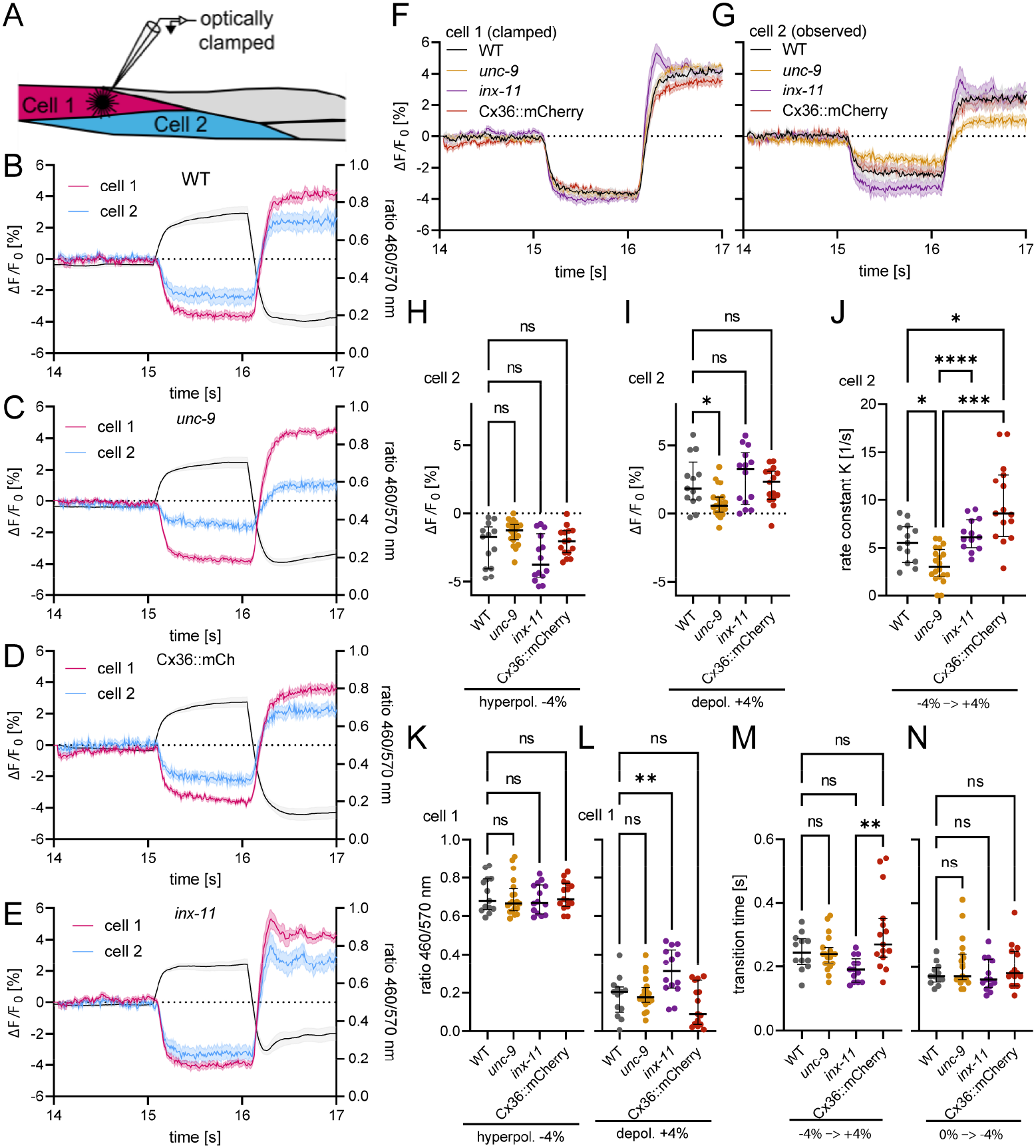
Cell-specific OVC measurements reveal reduced junctional conductance in *unc-9* mutants and increased excitability of *inx-11* mutants. (A) Schematic illustration of a cOVC measurement. Cell 1 is optically clamped to specific membrane potentials (i.e. ΔF/F_0_ fluorescence levels), while voltage fluorescence changes in cell 2 are observed only. Mean ΔF/F fluorescence (left y-axis) traces of clamped cell 1 and observed cell 2 and mean wavelength changes (right y-axis; represented as the ratio of light intensities of the respective wavelength) in cell 1 of WT animals (B), *unc-9* (C), Cx36::mCherry overexpressing muscles (D) and *inx-11* mutants (E). Number of animals n=13 (WT), 20 (*unc-9*), 15 (Cx36), 14 (*inx-11*). (F) Mean (±SEM) ΔF/F_0_ fluorescence of clamped cell 1 and unclamped cell 2 (G). (H) Changes of QuasAr2 fluorescence in cell 2 during hyperpolarization and (I) depolarization step. Median with interquartile range. Welch ANOVA with Dunnett’s test in H; one-way ANOVA with Sidak test in I (ns p>0.05, * p<0.05). (J) Calculated rate constant K of the −4% to +4% ΔF/F_0_ step was calculated using an exponential plateau function. Median with interquartile range. Welch ANOVA with Dunnett’s test (* p<0.05, *** p<0.001, **** p<0.0001). (K) Wavelength changes (ratio of intensities of light of 460 nm over 570 nm, applied to cell 1) during the hyperpolarization and (L) depolarization steps. Median with interquartile range. Welch ANOVA with Dunnett’s test in K; one-way ANOVA with Sidak test in L (ns p>0-05, ** p<0.01). (M) Transition times for 8% and 4% ΔF/F steps (N). Median with interquartile range. Kruskal-Wallis with Dunn’s test (ns p>0.05, ** p<0.01).

### The cell-specific cOVC reveals less GJ-mediated conductance in *unc-9*, and higher excitability in *inx-11* mutants

In WT animals, cell 1 was reliably clamped to holding values of −4 and +4% ΔF/F_0_, while changes of voltage-dependent fluorescence were detected in the neighboring cell 2 (Fig. 7B). The unclamped cell 2 of WT animals almost reached the same level of hyper- and depolarization as cell 1 with no obvious time delay. Also in *unc-9* mutants, cell 1 could be clamped to −/+4% ΔF/F_0_ (Fig. 7C, F). However, the observed cell 2 showed slightly reduced hyper-, and significantly reduced depolarization compared to WT cell 2 (Fig. 7C, G-I), indicating overall lower inter-cellular conductance. Transition from hyper-to depolarization occurred significantly slower in cell 2 of *unc-9* mutants compared to WT (Fig. 7J), indicating slower transmission of electrical signals, in agreement with the reported lower junctional conductance in *unc-9* mutants (12). Junctional coupling of Cx36::mCherry overexpressing muscles was comparable to WT: Cell 1 could be clamped to holding values of −/+4% ΔF/F_0_ and also the unclamped cell 2 reached similar levels as in WT during hyper- and depolarization (Fig. 7D, F-I). However, transition from hyper-to depolarization in cell 2 occurred significantly faster compared to WT (Fig. 7J), indicating faster transmission of electrical signals between adjacent cells, possibly due to an overall higher number of GJs. In the *inx-16* mutant background, expression of the QuasAr2 transgene was largely reduced, for unknown reasons. Last, we compared the junctional coupling of cOVC measurements in *inx-11* mutants. Clamping cell 1 of *inx-11* mutants was achievable, with cell 2 almost reaching the same voltage fluorescence values as cell 1 (Fig. 7E, F-I). Interestingly, cell 1 reached the same depolarization level as WT with less Chrimson activation (i.e., a higher blue-to-green ratio; 0.18±0.03 ratio 460/570 nm (WT) vs. 0.13±0.03 (*inx-11*)), while hyperpolarization was not affected (Fig. 7K, L). The transition time from the hyperpolarizing step to the depolarizing step was also significantly faster compared to WT (Fig. 7M). This indicates that the muscles of *inx-11* mutants exhibit increased excitability, hinting towards homeostatic changes of the other innexins. Alternatively, a lower overall amount of GJs causes single muscle cells to be more ‘isolated’, thus leading to decreased electrical coupling and more excitability of single cells. This may be in line with the finding that APs in *inx-11* mutants had a larger area under the curve and were longer lasting (FWHM was larger than in WT; Fig. 6F, G), an effect that could be expected if current dissipates less readily from single muscle cells. This might also explain the higher crawling speed we observed in *inx-11* mutants (see Fig. 1).

## Discussion

GJs play an important role in the function of multicellular organisms, as they are involved in the exchange of small metabolites, and also mediate direct electrical signaling between adjacent cells. New methods to study GJ-mediated coupling are required to better understand their function and the influence of their compositions on their function *in vivo*. Here, we established an all-optical approach to study GJ-mediated electrical coupling in live *C. elegans*. Using QuasAr2, we could investigate intrinsic BWM cell activity in immobilized animals in a non-invasive manner. This is advantageous to electrophysiology in dissected worms, where artificial buffers are used, possibly affecting GJ function. In addition, spontaneous intrinsic muscle activity can be assessed, giving new insights into possible alterations in mutants. Asynchrony of APs in neighboring cells can be detected with voltage imaging, as was previously shown by calcium imaging (42), however, voltage imaging directly assesses electrical activity that precedes Ca^2+^ fluctuations, and achieves higher temporal resolution. While voltage imaging allows the investigation of electrical coupling mediated by GJs between adjacent cells, other functions of GJs, like metabolite exchange, cannot be directly investigated using this approach.

We expanded the previously established OVC to allow optically clamping one specific cell and analyzing the induced membrane potential changes in the neighboring cells. With this cOVC, we could observe changes in GJ-mediated conductance in *unc-9* mutants and also increased excitability in the *inx-11* mutants. Like in the original OVC, changes in membrane potential are measured as the change of fluorescence. These values have been calibrated to actual voltages (38), thus enabling a realistic estimation from imaging data. The major benefit of the cOVC is that there are no non-native conditions as imposed by dissection and use of artificial intra- and extracellular solutions, as required for electrophysiology. Moreover, for technical reasons, electrophysiology recordings are mostly limited to BWMs between the neck and vulva, while voltage imaging can be conducted in any BWM cell, provided the expression of the voltage indicator is robust. This approach can now be expanded to neurons or other cell types. GEVIs are available across a wide spectral range (31, 36, 50), giving the possibility to combine different sensors and to establish dual-color voltage imaging in two different cell types. Additionally, the cOVC we established should be adaptable for the use in other cell types or even in other organisms, as was demonstrated for the original OVC (38). Due to different cellular properties, adaptations depending on the model organism must be taken into account. Regarding the investigation of GJ composition, the cOVC could be used to screen different combinations of innexins, which could be overexpressed in BWMs, as an amenable test system that provides also behavioral readouts.

Our functional analyses revealed that *unc-9* mutants showed overall lower correlation of neighboring muscle cells during spontaneous activity and decreased electrical coupling between adjacent cells. These findings are comparable to measured transjunctional currents, that were significantly decreased in *unc-9* mutants (12, 23, 42). Furthermore, even during spontaneous BWM activity, overall lower correlation of muscle cells was observed. Compared to the effects of the other investigated innexin mutants, loss of UNC-9 had the most prominent effect, indicating a major role of UNC-9 in muscular coupling. Yet, since not all activity is abolished in *unc-9* mutants, additional innexins play a role, and we observed functions of INX-11 and INX-16, just as previously reported (12). Interestingly, we did not observe increased excitability or input resistance in *unc-9* mutants, which would be expected when reducing leak currents to neighboring cells. Possibly, the presumable partner innexin INX-18 gets inserted into the membrane, and could form hemichannels. This insertion of hemichannels could maintain input resistance and excitability, while leading to reduced voltage signals to neighboring muscles. A similar hypothesis was raised by Güldenagel et al. to explain the observed decreased sensitivity in ON cone bipolar cells of Cx36-deficient mice (51). Hemichannel formation of *C. elegans* innexins was previously shown for UNC-7, where they function in thermosensory neurons to transmit temperature information (52). Also, UNC-9 mediates the spatial arrangement of GABAergic synapses in *C. elegans* in an (intercellular) channel-independent way (53).

In *inx-16* mutants we observed a higher peak amplitude and area under the curve in voltage imaging and electrophysiology experiments, as well as an increased input resistance, indicating a higher excitability of single muscle cells. This might result from overall fewer GJs and therefore less leak currents to neighboring cells. Alternatively, GJs that form in the absence of INX-16 exhibit different conduction properties. Less coupling synchrony between muscle cells was not observed in *inx-16* mutants, on the contrary, the cross-correlation of voltage signals in individual cells was significantly longer lasting, likely due to the broader AP burst episodes. *inx-11* mutants did not show any differences to WT in voltage imaging of spontaneous muscle activity. However, in patch-clamp recordings, broader APs were observed, and cOVC clamping of cell 1 required less Chrimson activation to achieve the same level of depolarization as WT, indicating that *inx-11* mutants also exhibit increased excitability. Interestingly, *inx-16* and *inx-11* mutants showed different crawling behaviors. While in *inx-16* mutants crawling was slightly impaired, *inx-11* mutants showed increased basal crawling speed. This could imply that these two innexins do not belong to the same population of GJs or that homeostatic changes in either the expression or localization of the remaining innexins occur. To achieve a more detailed understanding, the exact composition of GJs in BWMs needs to be determined, as well as the composition resulting from possible compensatory changes in the absence of certain innexins. Previous studies showed that the localization of innexins assembling into a distinct GJ depends on the localization of the partner innexins (54). In *Drosophila*, it was shown that heteromerization of innexin2 and 3 was essential for the correct localization of these two innexins. Deletion of one partner led to mislocalization to the cytoplasm, causing cell polarity defects (55). In *C. elegans*, heterotypic GJs consisting of UNC-9 and UNC-7S are formed between the AVB neuron and motor neurons, and the correct localization of the two innexins was shown to be interdependent (16). In addition, the deletion of one subunit of a heterotypic or heteromeric GJ could lead to changes in conductivity, e.g. by formation of GJs of different composition. That altered GJ composition can change conductivity was previously observed for UNC-9 and UNC-7 containing GJs, where heterotypic channels exhibit rectification, while homomeric channels do not (16, 56).

Overexpression of heterologous GJs, i.e. murine Cx36, led to increased correlation between muscle cells, and a faster transition from hyper-to depolarization in cOVC measurements, indicating more efficient and faster transmission of electrical signals between cells. This might be due to an overall increased junctional conductance, resulting in currents flowing more efficiently between neighboring cells. Since Cx36 GJs exhibit low sensitivity to transjunctional voltage and membrane voltage, this would allow the Cx36 channels to be open between cells undergoing strong fluctuations in their membrane potential (57). This increased conductance caused a severe disruption of normal locomotion behavior. Overexpression may additionally have caused localization of GJs/GJ plaques to regions of the BWMs that are not normally electrically connected, resulting in further disruption or interference of normal progression of excitation and thus the undulatory wave of muscle contraction. This indicates that muscular GJs indeed influence wave propagation, by a finely tuned synchronization of individual muscle activity. Possibly, the sequence of motor neuron activation needs to precisely time offsets in muscle activation, which become ‘smeared out’ on the level of individual muscle cells if the ensemble of GJs is not properly formed. Overall, our findings indicate that knockout of certain GJ subunits influences electrical signaling to a different extent, depending on the innexin deleted. *unc-9* mutants show the most prominent effect when looking at intrinsic muscle activity, with decreased correlation and desynchronized APs, suggesting a major role of UNC-9 in mediating the signal propagation in muscle cells. The effect of the loss of INX-11 and INX-16 was less prominent. The higher amplitude and area under the curve of *inx-16* APs, detected in both electrophysiology recordings and voltage imaging experiments, as well as the higher input resistance, suggest a higher excitability of single muscle cells, and further, that GJs actually may be involved in shaping the AP waveform. This might be a result of lower leak currents due to a lower number of GJs, as also indicated by the much larger voltage change in response to increasing current steps. Our work shows that network dynamics, also in the BWM ensemble, depend on the right balance of GJs. Reduced coupling results in loss of synchrony, while increased coupling leads to an over-synchronized state that reduces dynamic flexibility and spatially diverse activity of individual cells within the BWM ensemble.

## Materials and methods

### Molecular biology

Plasmid pAB16 (*pmyo-3::QuasAr*; Addgene plasmid #130272) was previously described (36).

**pAB13 (*pmyo-3::ChR2(H134R)::mTFP*)** was generated via subcloning of mTFP (pFR4545, gift from Stefan Eimer) into *pmyo-3::ChR2(H134R-g*.*o*.*f)::YFP*, using restriction enzymes *NotI* and *EcoRI*.

**pCR03 (*pmyo-3::Cx36::mCherry*)** was generated by amplification of Cx36 sequence with primers oCW19 (5’ AAGACTGGAGAAGTCACTGctagATGGGAGAGTGGACCATCCTC 3’) and oCW20 (5’ CGCCCTTGCTCACCATCTCGAgGACGTAGGCGGAGTCGGAGGATTG 3’) and cloning it into the vector pED031 (*pmyo-3::CD4-2::mCherry*) using *XhoI* and *NheI* and Gibson Assembly.

*C. elegans* strains

*Caenorhabditis elegans* strains were cultivated on nematode growth media (NGM) plates seeded with *Escherichia coli OP50-1* strain (Brenner, 1974). Strains used in this study are listed in Table 1. Transgenic strains were generated by microinjection of plasmid DNA into the gonads of either wildtype (Bristol N2) or *juSi164* (Histone-miniSOG) animals (Mello et al., 1991)(58). Integration of transgenes was performed as previously described (Dunkel et al., 2025; Noma and Jin, 2018), strains were then outcrossed to wild type N2.

**Table 1:** *C. elegans* strains used in this study.

| Strains | Genotype | Source |
| --- | --- | --- |
| N2 | *C. elegans* wildtype isolate | CGC |
| CB101 | *unc-9(e101)X* | CGC |
| RB2108 | *inx-11(ok2783)V* | CGC |
| ZX3231 | *inx-16(tm1589)I* | NBRP |
| ZX2476 | *zxEx1139[pmyo-3::QuasAr; pmyo-2::CFP]* | (36) |
| ZX2876 | *zxIs139[pmyo-3::GtACR2::mCerulean::βHK::Chrimson; pmyo-3::QuasAr; pmyo-2::CFP]* | (38) |
| ZX3194 | *unc-9(e101)X; zxEx1139[pmyo-3::QuasAr; pmyo-2::CFP]* | this study |
| ZX3225 | *inx-11(ok2783)V; zxEx1384[pmyo-3::ChR2(H134R)::mTFP; pmyo-3::QuasAr; pmyo-2::CFP]* | this study |
| ZX3226 | *inx-16(tm1589)I; zxEx1384[pmyo-3::ChR2(H134R)::mTFP; pmyo-3::QuasAr; pmyo-2::CFP]* | this study |
| ZX3227 | *zxEx1384[pmyo-3::ChR2(H134R)::mTFP; pmyo-3::QuasAr; pmyo-2::CFP]* | this study |
| ZX3479 | *zxls190[pmyo-3::QuasAr; pmyo-2::CFP]; zxls5[punc-17::ChR2(H134R)::yfp; lin-15+]* | this study |
| ZX4068 | *unc-9(e101)X; zxls139[pmyo-3::GtACR2::mCerulean::βHK::Chrimson; pmyo-3::QuasAr; pmyo-2::CFP]* | this study |
| ZX4095 | *inx-11(ok2783)V; zxls139[pmyo-3::GtACR2::mCerulean::βHK::Chrimson; pmyo-3::QuasAr; pmyo-2::CFP]* | this study |
| ZX4101 | *zxls305[pmyo-3::Cx36::mCherry, pmyo-2::CFP]* | this study |
| ZX4102 | *zxls190[pmyo-3::QuasAr; pmyo-2::CFP]; zxls5[punc-17::ChR2(H134R)::yfp; lin-15+]; zxls305[pmyo-3::Cx36::mCherry, pmyo-2::CFP]* | this study |
| ZX4103 | *zxls139[pmyo-3::GtACR2::mCerulean::βHK::Chrimson; pmyo-3::QuasAr; pmyo-2::CFP]; zxls305[pmyo-3::Cx36::mCherry, pmyo-2::mCherry]* | this study |

### Behavioral assays

For investigation of swimming behavior, L4 larvae were picked onto newly seeded NGM plates. After 12-16 hours, the animals were washed three times with M9 buffer and transferred using 800 µl M9 buffer to unseeded NGM plates (3.5 cm diameter). Animals were kept in the dark for 15 minutes to accommodate swimming behavior prior to the measurements. Swimming behavior was recorded using the multiworm tracker (MWT) setup (59), equipped with a camera (Falcon 4M30, DALSA) and an infrared transmission light source (6 WEPIR3-S1, Winger, 850 nm, 3W). Videos of the swimming behavior were recorded using a LabVIEW-based custom software (MS-Acqu) and were analyzed as previously described (60).

For crawling behavior, preparation was the same as for swimming assays, however, the animals were transferred in a droplet of M9 buffer to unseeded NGM plates (6 cm diameter). Crawling behavior was recorded using the MWT setup. The software ‘Multiworm tracker’ (version 1.3) was used to record speed data and tracks were extracted using Choreography (version 4.2) (59).

### Fluorescence imaging

Animals were placed onto agarose pads containing 10% agarose dissolved in M9 buffer and immobilized using 0.1 µm PolyBeads (at 2.5% w/v, Polysciences). Images were acquired on an inverted microscope (Zeiss Axio Observer Z1) equipped with a sCMOS camera (Kinetix22, Teledyne) and a 40x oil immersion objective (Zeiss EC Plan-NEOFLUAR x40/N.A1.3, Oil DIC ∞/0.17). To obtain representative fluorescent images, mCherry was excited with a 590 nm LED (Lumen 100, Prior Scientific) and using a mCherry filterset (mCherry HC, AHF Analysentechnik).

### Voltage imaging of spontaneous muscle cell activity

For voltage imaging of spontaneous muscle cell activity, L4 larvae were picked onto NGM plates seeded with OP50 suspension containing all-*trans* retinal (ATR) (stock in ethanol, final concentration: 0.05 mM). Animals were kept in the dark overnight. For imaging, animals were placed onto pads containing 10% agarose dissolved in M9 buffer and immobilized using 0.1 µm PolyBeads. Imaging was performed on an inverted microscope (Zeiss Axio Observer Z1), equipped with a 40x oil immersion objective (Zeiss EC Plan-NEOFLUAR x40/N.A1.3, Oil DIC ∞/0.17), a laser beam splitter (HC BS R594 lamda/2 PV flat, AHF Analysentechnik), a Galilean beam expander (BE02-05-A, Thorlabs), and a sCMOS camera (Kinetix22, Teledyne). QuasAr2 fluorescence was excited with a 637 nm red laser (OBIS FP 637LX, Coherent) at 1.8 W/mm^2^ and imaged at 700 nm (700/75 ET Bandpass filter, integrated in Cy5 filter cube, AHF Analysentechnik). Voltage imaging of spontaneous BWM activity of *inx-11, −16* and Cx36 overexpressing strains was performed at around 100 fps, with 10 ms exposure time and 4×4 binning. Voltage imaging of *unc-9* mutants was performed at around 20 fps. Videos were analyzed in MicroManager 1.4 using a custom-written bleach correction script (‘offline_bleach_correction.bsh’). For this, a region of interest (ROI) was set around a specific muscle cell and a background ROI was defined inside the worm. Fluorescence was bleach- and background-corrected, and the fluorescence change rate (ΔF/F) was calculated. The bleach-corrected values for the fluorescence change rate (ΔF/F) were used for analysis. Since during strain construction different QuasAr2 arrays had to be used, we compared the three WT controls for overall correlation of muscle activity, and found no significant differences (Supp. Fig. 1D).

Peak analysis was conducted using a custom-written MATLAB R2024A script (Mathworks, Natick, MA, USA). The baseline was interpolated from minima and the peaks were detected and extracted using the ‘findpeaks’ function. Subsequently, peak height, area and FWHM were calculated. Correlation of spontaneous muscle activity was calculated using Pearson correlation using GraphPad Prism (GraphPad Software Inc., Boston, MA, USA). Cross correlation was calculated using the ‘Xcorr’ function in MATLAB. For cross correlation analysis of single peaks, ΔF/F traces were imported, peaks were identified using the ‘findpeaks’ function and 0.25 s before and after the peaks were extracted. For every extracted peak, the ‘Xcorr’ function was run for all three combinations of cells (cells 1,2; 1,3; 2,3). For analysis, the mean of the first ten peaks and cross correlations were calculated. For cross correlations of longer time windows, ΔF/F traces were imported, peaks were identified using the ‘findpeaks’ function and 0.65 s before and after the peak were extracted. For every extracted peak, the ‘Xcorr’ function was run for the three combinations. For analysis, the mean of the first five cross correlations was calculated.

### cOVC measurements

For cOVC measurements, animals were supplemented with ATR, by picking L4 larvae onto NGM plates containing OP50 mixed with ATR (dissolved in ethanol, final concentration: 0.02 mM). Animals were kept in the dark overnight. Animals were immobilized in 0.1 µm PolyBeads on 10% agarose pads. Voltage-dependent QuasAr2 fluorescence was imaged with a 637 nm red laser (OBIS FP 637LX, Coherent) equipped with a Galilean beam expander (BE02-05-A, Thorlabs) at 1.8 W/mm^2^ and imaged at 700 nm (700/75 ET bandpass filter, integrated in Cy5 filter cube, AHF Analysentechnik). BiPOLES constituents were activated using an Optoma UHD38x projector (lamp DLP, expansion lens removed according to (61). Projector light was cleaned up by a 1% transmission ND filter, a GFP/mCherry ET dual band excitation filter (AHF Analysentechnik), and a single notch filter for 488 nm (AHF Analysentechnik). Imaging was performed on a Zeiss Axiovert 200 inverted microscope equipped with a laser beam splitter (HC BS R594 lambda/2 PV flat, AHF Analysentechnik), 40x oil immersion objective (Zeiss EC Plan-NEOFLUAR x40/N.A1.3, Oil DIC ∞/0.17) and sCMOS camera (Kinetix22, Teledyne). The complete setup is shown in Supp. Fig. 3A. All OVC experiments were performed at 100 fps with 10 ms exposure, a binning of 4×4 and the camera’s sensor readout was reduced to a 350×350 px region. The pixel offset, i.e. the difference of x and y coordinates between a set ROI and the camera image was 1.3 ± 0.3 px / 0.91 ± 0.21 µm, where 50 µm corresponded to 77 px. The light intensity measured at the focal plane of the sample was 120 µW/mm^2^. The time lag between a command of the software (see below) to send an illumination pattern and the observed camera signal was 56.1 ± 2.3 ms (@ 100 fps acquisition of the Kinetix22 camera in dynamic range); the field of view was set to 350×350 px; the projector was running at 240 Hz image refresh rate and 800×600 pixel resolution.

The specifications, parameters and algorithm were previously described (38), and adapted accordingly in the following. For cOVC measurements, no Kalman-filter was applied for sensor-signal smoothing. The cOVC control software was written in Beanshell and initiated *via* the µManager script panel. The microscope setup uses a projector whose output must be calibrated relative to the imaging system (‘projector_calibration.bsh’). Characterization of the illumination system was performed according to (61). For this purpose, a calibration pattern is projected consisting of five adjustable circles (size, color, and relative spacing). By manually localizing the centers of these circles in the camera image, the experimenter can determine lateral x- and y-offsets as well as the magnification factor between the projected pattern and the recorded field of view. These parameters are then applied to correct the projection in subsequent experiments. This procedure ensures that the projected ROIs precisely match with the outlines of the targeted cells, minimizing unintended activation of neighboring cells. After the calibration circles have been brought into focus both at the focal plane and at the outer edges of the field of view (to ensure full FOV coverage), a snapshot is acquired. This image serves as the basis for the manual localization of the circle center coordinates. The software allows the image to be flipped horizontally and/or vertically for calibration, if required by the respective camera-microscope configuration. After the circle centers have been manually selected, the software computes the lateral x- and y-offsets as well as the magnification factor between the projected pattern and the acquireded camera snapshot. These calibration parameters are then stored in an output file for use as correction parameters in the subsequent cOVC experiments.

Prior to running the cOVC script (‘Multicell_OVC.bsh’), the user needs to acquire an image of the cells to be measured. An input tab allows the user to select the acquisition parameters. The number of frames and the holding values can be selected for each step. Cell 2 can be clamped or observed only. Further, the number of frames for the calibration phase, tolerance range (in %) and the increment in which the control variable will be adapted at each point can be changed. The control variable is the color ratio between the blue and green channel and can be set between 0 (full green) and 1 (full blue). The ratio is computed from the underlying RGB intensity values of the blue and green channels, which range from 0 to 255. It will change at each step by 0.005 weighted by the control error, which is the deviation between actual gray value and desired holding value. The user can also adapt the multiplier strength, which is the strength in which the control variable is adjusted, as well as the range of available color ratios, by changing the upper and lower limits and the calibration start color ratio. The latter is the color presented during the calibration phase to counteract pre-activation of Chrimson by the 637 nm laser for QuasAr2 illumination. If it is intended to only observed cell 2, its initial color ratio is kept constant throughout the experiment to prevent unwanted Chrimson activation. Next, by following instructions, ROIs for cell 1 and 2, as well as a background ROI are selected. The script prompts the user to turn on the 637 nm laser and, after pressing ‘ok’, the acquisition is running. Preceding the clamping phase, there is a calibration phase in which the acquired gray values are used to evaluate the parameters for the exponential decay function to correct for photobleaching of QuasAr2. Once the OVC acquisition is complete, a results text file and an image stack file are saved automatically.

### Electrophysiology

Electrophysiology recordings of BWM cells were done in dissected adult worms as previously described (62). Animals were immobilized with Histoacryl L glue (B. Braun Surgical, Spain) and a lateral incision was made to access NMJs along the anterior ventral nerve cord. The basement membrane overlying BWMs was enzymatically removed by 0.5 mg/ml collagenase for 10 s (C5138, Sigma-Aldrich, Germany). Integrity of BWMs and nerve cord was visually examined via DIC microscopy. Recordings from BWMs were acquired in whole-cell patch-clamp mode at 20−22 °C using an EPC-10 amplifier equipped with Patchmaster software (HEKA, Germany). The head stage was connected to a standard HEKA pipette holder for fire-polished borosilicate pipettes (1B100F-4, Worcester Polytechnic Institute, USA) of 4–10 MΩ resistance. The extracellular bath solution (CRG) consisted of 150 mM NaCl, 5 mM KCl, 5 mM CaCl_2_,1 mM MgCl_2_,10 mM glucose, 5 mM sucrose, and 15 mM HEPES, pH 7.3 (adjusted with NaOH), ~330 mOsm. The internal patch pipette solution consisted of K-gluconate 115 mM, KCl 25 mM, CaCl_2_ 0.1 mM, MgCl_2_ 5 mM, BAPTA 1 mM, HEPES 10 mM, Na_2_ATP 5 mM, Na_2_GTP 0.5 mM, cAMP 0.5 mM, and cGMP 0.5 mM, pH 7.2 (adjusted with KOH), ~320 mOsm.

Membrane potential and APs in BWMs were recorded in current clamp mode. To induce APs, a current pulse of +20 pA (50 ms) was injected via the Patchmaster software. For data analysis, voltage traces were aligned to the peak of the AP, and replotted, thus appear shifted relative to the current pulse.

For measurements of input resistance (R_in_), a current pulse of −20 pA was injected via the Patchmaster software for 1000 ms in three consecutive pulses with 1000 ms breaks in between the current pulses. Alternatively, current pulses of −10, −20, and −30 pA were injected for 1000 ms. For normalized input resistance, R_in_ was divided by the membrane capacitance (C_m_) of the respective cell.

### Statistical analysis

Statistical analysis used is described in the figure legends. Data was tested for normality prior to statistical analysis. Statistical analyses were performed in GraphPad Prism (Version 9.4.1, GraphPad Software Inc., Boston, MA, USA).

## Acknowledgements and funding sources

We are indebted to Katharina Kuhlmeier and Franziska Baumbach for expert technical assistance and providing infrastructure, and to members of the lab for support and critical feedback. We thank William Schafer for the Cx36 plasmid. Some strains have been obtained from the *Caenorhabditis* Genetics Center, which is funded by NIH Office of Research Infrastructure Programs (P40 OD010440), and from the National Bioresource Project for the Experimental Animal “Nematode *C. elegans*”. This work was funded by the Deutsche Forschungsgemeinschaft (DFG), grant CRC1507, P06.

## Author contributions

Conceptualization: NE, AB, JL, AG.

Methodology: NE, CW, AB, JL.

Investigation: NE, CW, AB, JL.

Formal Analysis: NE, CW, AB, JL, AG.

Writing (Original Draft): NE, AB.

Writing (Review & Editing): NE, JL, AG.

Supervision: AG.

Funding Acquisition: AG.

## Supplementary Material

**Supplementary Figure 1:**
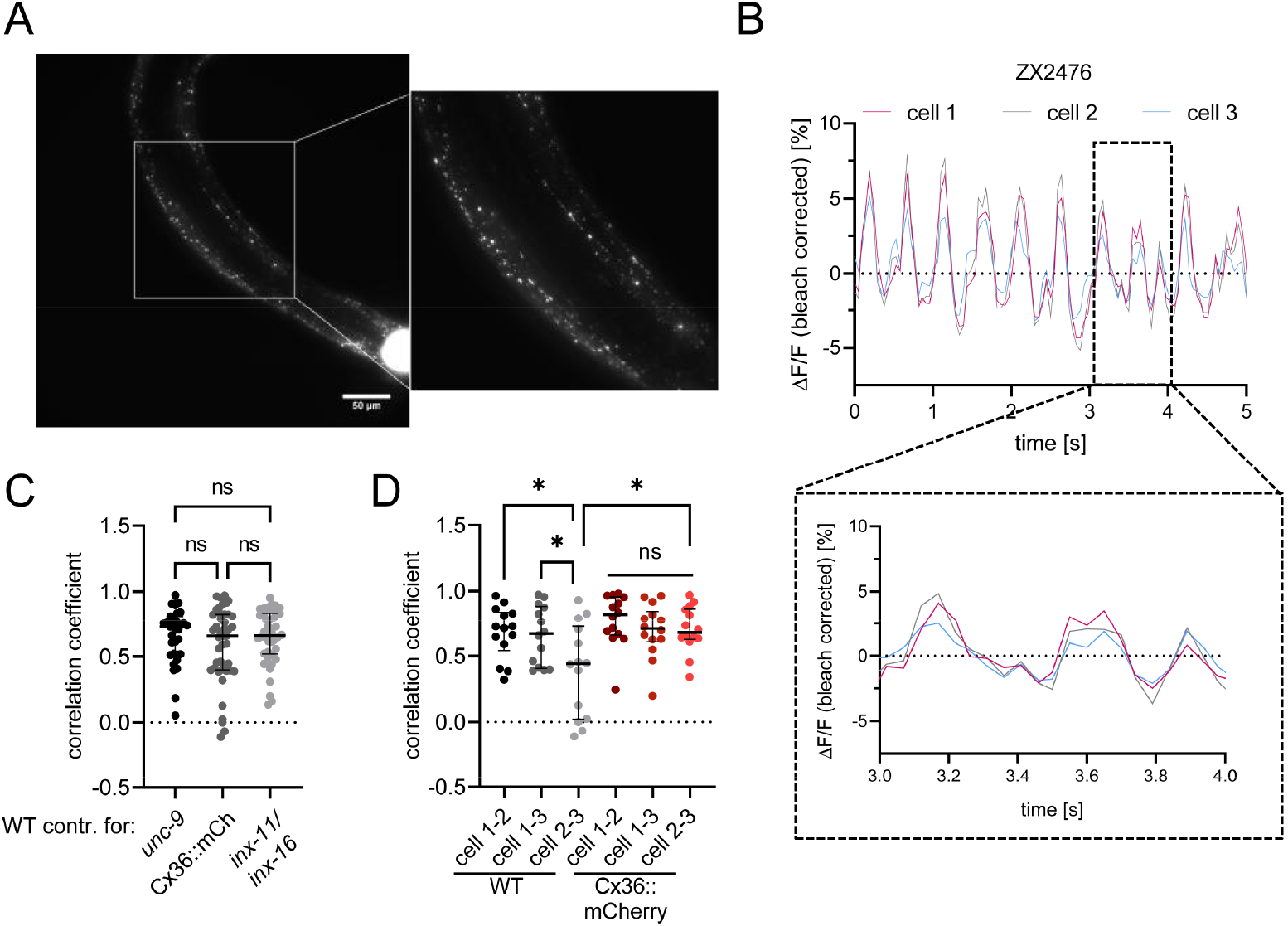
(A) Fluorescence micrograph showing expression pattern of Cx36::mCherry in BWMs. Scale bar: 50 µm. (B) Representative ΔF/F fluorescence time traces of strain ZX2476, the WT control for *unc-9* mutants. (C) Correlation coefficients of all three different WT strains expressing different QuasAr2 arrays were calculated using Pearson correlation. Median with interquartile range. Kruskal-Wallis with Dunn’s test (ns p>0.05). (D) Correlation coefficients for the three pairs of Cx36 overexpressing muscle cells analyzed in an ensemble were calculated using Pearson correlation. Median with interquartile range. Kruskal-Wallis with Dunn’s test (** p<0.01).

**Supplementary Figure 2:**
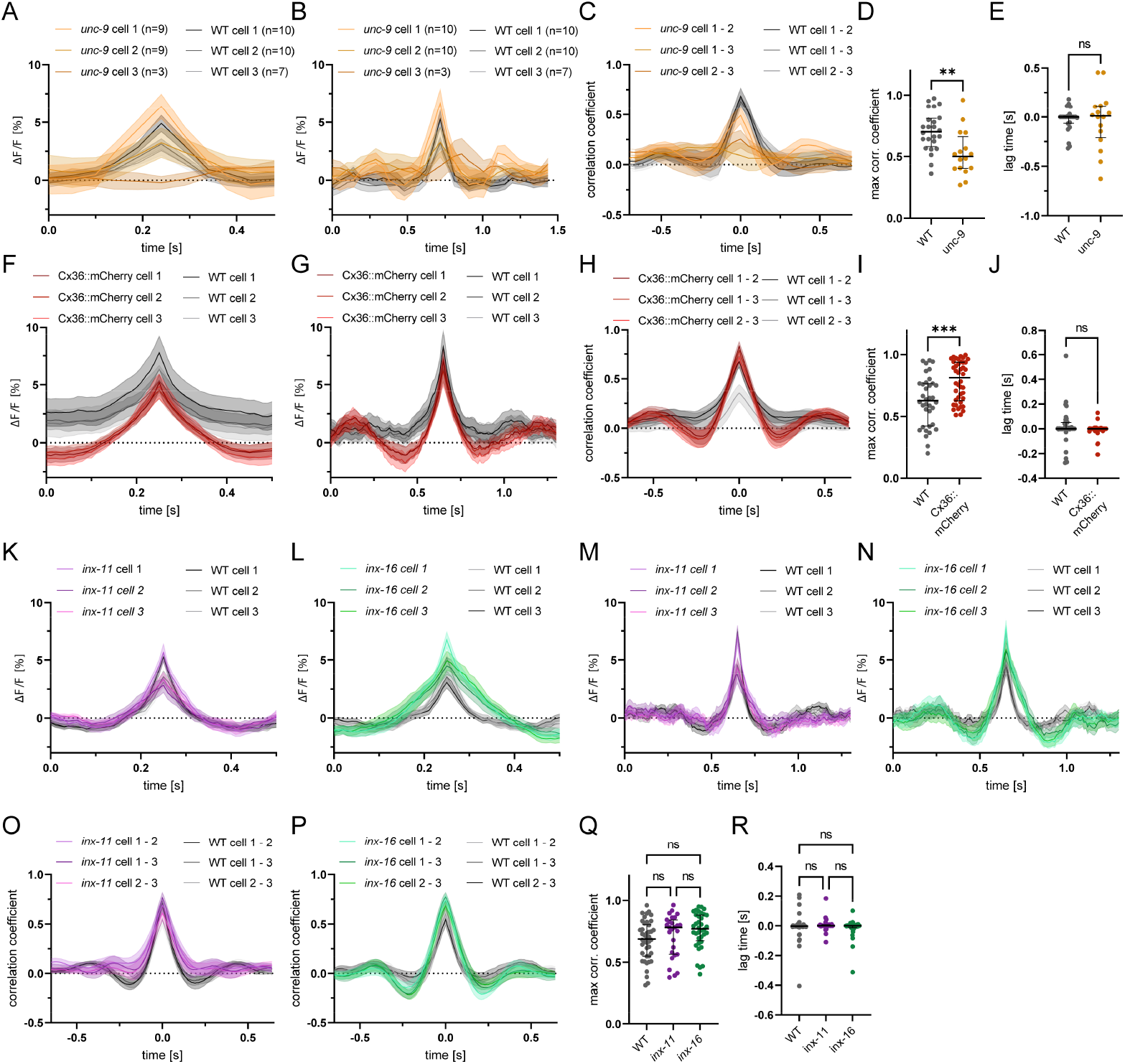
(A) Mean (±SEM) ΔF/F traces of extracted peaks (±0.25 s) used for cross correlation analysis of single peaks of *unc-9* mutants, (F) Cx36::mCherry, (K) *inx-11* and (L) *inx-16* animals, as well as their respective WT controls. (B) Mean (±SEM) ΔF/F traces of extracted peaks (±0.65 s) used for cross correlation analysis of longer time windows of *unc-9* mutants, (G) Cx36::mCherry, (M) *inx-11* and (N) *inx-16*, as well as their respective WT controls. Number of animals n=7-10 (WT), 3-10 (*unc-9*) in B, n=12 (WT), 14 (Cx36) in G, n=14 (WT), 9 (*inx-11*), 12 (*inx-16*) in K and L. (C) Cross-correlation analysis of longer time windows (±0.65 from peaks) of pairs of the three-cell ensemble of *unc-9* mutants, (H) Cx36::mCherry animals, (O) *inx-11* and (P) *inx-16* mutants. (D) Maximum correlation coefficient of *unc-9*, (I) Cx36 animals, and (Q) *inx-11* and *inx-16* mutants of the cross correlations of five extracted time windows from each animal. (E) Lag time at maximum correlation of *unc-9*, (J) Cx36 animals, and (R) *inx-11* and *inx-16* mutants of the cross correlations of five extracted time windows from each animal. Each dot represents the mean of the first 5 cross correlations of one pair of muscle cells. Median with interquartile range. Unpaired t test in D, Welch t test in E, Mann-Whitney test in I and J, Kruskal-Wallis with Dunn’s test in Q and R (ns p>0.05, ** p<0.01, *** p<0.001).

**Supplementary Figure 3:**
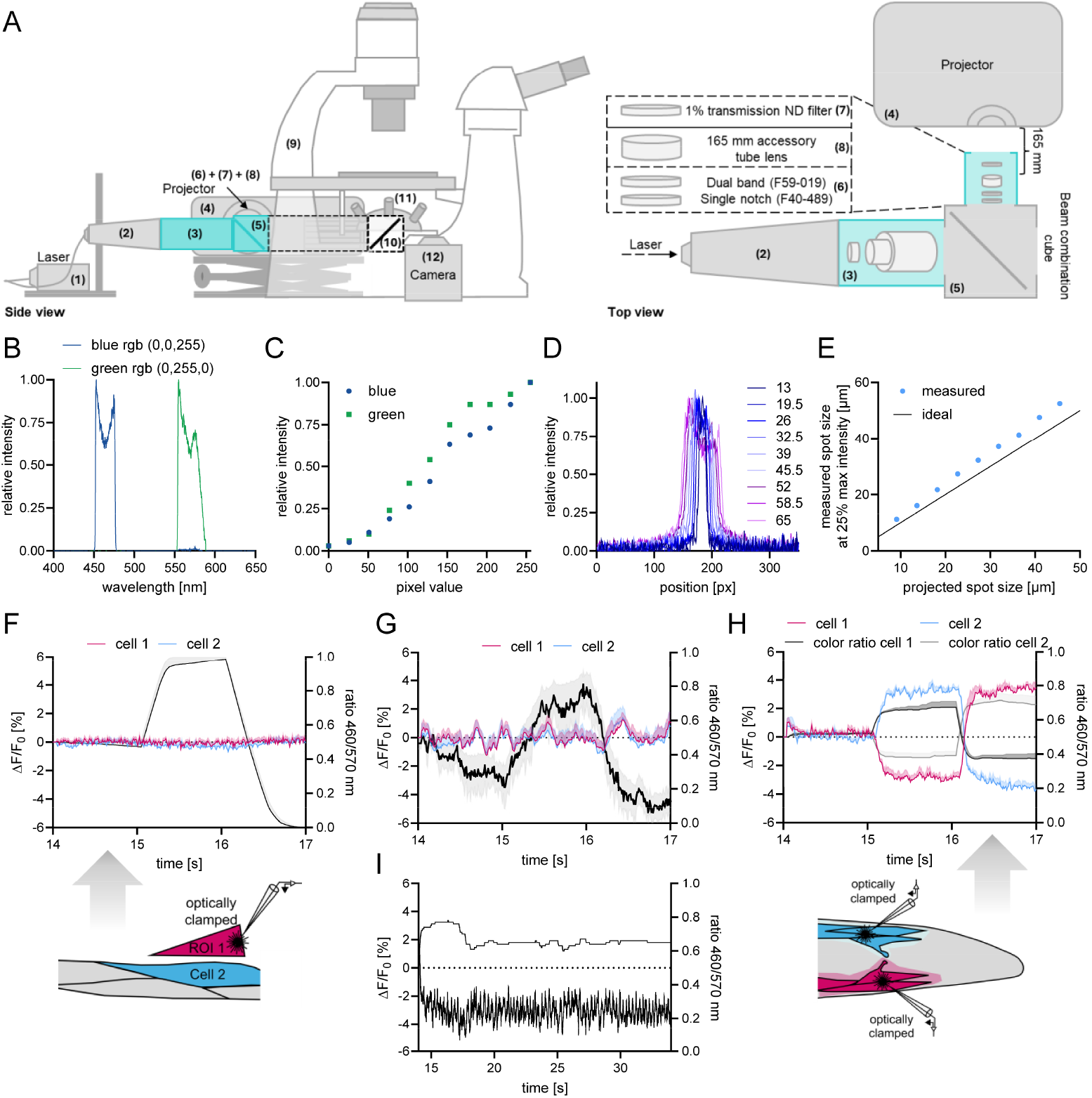
(A) Hardware setup for cOVC measurements: QuasAr2 is excited with a 637 nm laser [1], equipped with a Galilean beam expander [2] at 1.8 W/mm^2^ and imaged at 700 nm, using a 700/75 ET bandpass filter [10]. BiPOLES components were activated using light from a video projector with digital light processing unit [4]. Projector light was narrowed by additional filters [6]. In front of the accessory tube lens [8] a 1% transmission ND filter [7] was added. Between the beam combination tube equipped with a laser beam splitter [5], and the Galilean beam expander [2], an adapter with lenses of the epifluorescence path of the microscope [3] was added. Imaging was performed on a Zeiss Axiovert 200 inverted microscope [9] equipped with a 40x oil immersion objective [11] and a Kinetix22 sCMOS camera [12]. (B) Spectral characteristics of beamer output were shaped with excitation filters and ND filters. (C) Relative intensity as a function of color pixel value (0 (full off) – 255 (full on)) for blue and green RGB colors. (D) Measured width of different projected spot sizes using a 40x objective. (E) Measured spot sizes using a 40x objective. Width was measured from traces shown in (D) at the point where intensity drops to 25% of maximum value.. (F) cOVC measurement, where the ROI for ‘clamped cell 1’ was moved outside of the worm, adjacent to a BWM cell (ROI 2). Illumination protocol of ROI1, going from equal blue and green illumination to 100% blue and then to 100% green, is indicated (black curve, right y-axis; N=13 experiments). (G) Mean±SEM ΔF/F_0_ traces of strain only expressing QuasAr2 in BWMs (red and blue traces for cells 1 and 2), i.e., without BiPOLES, while the cOVC stimulation program attempts to run a 0, −4, +4% ΔF/F protocol (black traces, mean±SEM). N=15 experiments were averaged. (H) Independent clamping of individual BWMs located at opposite muscle strands in the head; mean±SEM ΔF/F_0_ traces of QuasAr2 fluorescence of the two muscles, indicated in blue and red color. The light stimulation protocol is indicated in black and grey for the opposite cells, as indicated. Average of N=5 experiments. (I) Long term −3% ΔF/F_0_ clamping of one individual cell. Shown are the achieved fluorescence values (black trace, bottom) and the ratio of the required wavelengths illuminated on the cell (upper half, referring to right y-axis).

## Supplementary Movies

**Supplementary Movie 1:** Three muscle cells expressing QuasAr2, fluorescence is imaged. Fluctuations in fluorescence intensity result from spontaneous membrane voltage changes.

**Supplementary Movie 2:** Example of an cOVC experiment. Two neighboring muscle cells in the center are outlined with regions of interest, labeled cell 1 and cell 2. Initially, a 15 s calibration period (indicated by the label and progressing line from left to right) is used to record fluorescence and photobleaching of QuasAr2, in order to calculate correction parameters for the clamping phase. Calibration is followed by consecutive steps of clamping voltage-dependent fluorescence of cell 1 to 0% ΔF/F_0_, −4%, and +4%, indicated by the downward and upward deflected progressing line at the bottom of the image.

